# The *Petasites hybridus* CO_2_-extract (Ze 339) blocks SARS-CoV-2 replication *in vitro*

**DOI:** 10.1101/2021.12.03.471068

**Authors:** Lorena Urda, Matthias Heinrich Kreuter, Jürgen Drewe, Georg Boonen, Veronika Butterweck, Thomas Klimkait

## Abstract

The coronavirus disease 2019 (COVID-19), caused by a novel coronavirus (SARS-CoV-2), has spread worldwide, affecting over 250 million people and resulting in over five million deaths. Antivirals that are effective are still limited. The antiviral activities of the *Petasites hybdridus* CO_2_-extract Ze 339 were previously reported. Thus, to assess the anti-SARS-CoV-2 activity of Ze 339 as well as isopetasin and neopetasin as major active compounds, a CPE- and plaque reduction assay in Vero E6 cells was used for viral output. Antiviral effects were tested using the original virus (Wuhan) and the Delta variant of SARS-CoV-2. The antiviral drug remdesivir was used as control. Pre-treatment with Ze 339 in SARS-CoV-2 infected Vero E6 cells with either virus variant significantly inhibited virus replication with IC_50_ values of 0.10 and 0.40 μg/mL, repectively. The IC_50_ values obtained for isopetasin ranged between 0.37-0.88 μM for both virus variants, that of remdesivir between 1.53-2.37 μM. In conclusion, Ze 339 as well as the petasins potently inhibited SARS-Cov-2 replication *in vitro* of the Wuhan and Delta variants. Since time is of essence in finding effective treatments, clinical studies will have to demonstrate if Ze339 can become a therapeutic option to treat SARS-CoV-2 infections.

## 1. Introduction

COVID-19 (<u>co</u>rona<u>vi</u>rus <u>d</u>isease 20<u>19</u>) first occurred in China end of 2019 and has since then spread around the globe, causing more than 5 million deaths [1]. Severe acute respiratory syndrome coronavirus-2 (SARS-CoV-2) is transmitted mainly via droplets and aerosols (for review see [2]). Infectivity begins one to several days before symptom onset, and even asymptomatically infected individuals can transmit the virus [2]. The disease affects the upper respiratory tract and lungs, heart, liver, gastrointestinal tract, and other organs, but the majority of infections remains asymptomatic, and patients develop only mild symptoms [3]. However, severe courses of COVID-19 are commonly accompanied by severe immune activation [4]. Observations include lymphocytopenia, an increase in neutrophils, and increased serum levels of IL-1β, IL-2, IL-4, IL-6, IL-10, TNF-α, and interferon-γ [3, 5, 6]. This cytokine response (also referred to as ‘cytokine storm’) is associated with cellular injury, which in turn is reflected in increased serum levels of lactate dehydrogenase (LDH), cardiac and hepatic enzymes, and the activation of coagulation and fibrinolysis with markedly increased plasma levels of D-dimers, among others [3, 5, 6]. In acute respiratory distress syndrome (ARDS) due to COVID-19, alveolar injury with desquamation of pneumocytes, hyaline membranes, and lymphomonocytic infiltrates has been described [7-9]. Lymphocytic endothelitis and apoptosis of endothelial cells in lungs, kidneys, and small intestine have also been observed [10]. The hyperinflammatory host-response caused by an overproduction of early response proinflammatory cytokines (e.g. TNFα, IL-6, IL1-ß) leads to multiorgan thrombotic complications, hyperpermeability, organ failure and death [11, 12]. For severe cases, several therapeutic strategies are currently being explored targeting the hyperinflammation caused by an overactive cytokine response (for review see [13]). However, a breakthrough in drug therapy has not yet been achieved, although the first candidates for an oral therapy have reached the approval process [14]. Among those is Paxlovid™ with the active ingredient PF-07321332 which is the first oral antiviral drug available desigend to combat SARS-CoV-2. PF-07321332 is an antiviral medication developed by Pfizer that inhibits the activity of 3C-like protease (3CL^PRO^) required for virus replication. Without the activity of the SARS-CoV-2 3CL^PRO^, nonstructural proteins (including proteases) cannot be released to perform their functions, thereby inhibiting viral replication [15].Paxlovid™ also contains a modest dose of ritonavir (a Ser-protease inhibitor and inhibitor of cytochrome P450 3A4), which delays the breakdown of PF-07321332, allowing it to stay in the body for longer at virus-inhibiting levels. In COVID-19 patients, the drug is likely to lower the rates of hospitalization [16]. Another drug currently under investigation is Molnupiravir, a Ridgeback Biotherapeutics product licensed by Merck, which increases the frequency of spontaneous alterations in the viral RNA and thus, inhibits SARS-CoV-2 replication [14]. In the numerous clinical trials that are currently being conducted worldwide, only remdesivir and dexamethasone have shown partial efficacy in combating hyperinflammatory stages of severe COVID-19 [17-24]. Furthermore, monoclonal antibodies, which block coronavirus surface proteins and reduce SARS-CoV-2 infection, are the only type of therapy approved in the United States for early-stage COVID-19. However, the high cost (about $2000 per dose, currently subsidized by the federal government in the United States), scarcity, and necessity to infuse or inject them have hindered their usage, especially in the developing world [14]. Therefore, COVID-19 therapies that are both affordable and simple to use are critically needed. Recent studies suggest that in addition to cytokines, leukotrienes (LTs) could also be considered as therapeutic targets since they further contribute to the hyperinflammation in severe COVID-19 cases [25-29]. The involvement of LTs in severe COVID-19 cases has been shown in some recent studies, confirming high LT levels in the bronchoalveolar lavage fluid of COVID-19 patients [30, 31].

Leukotrienes (LTB_4_, LTC_4_, LTD_4_, and LTE_4_) are peptide-conjugated lipids that are prominent products of activated eosinophils, basophils, mast cells and macrophages [32-34]. They are generated *de novo* from cell membrane phospholipid-associated arachidonic acid via the 5-lipoxygenase pathway. Known to cause contraction of bronchial smooth muscle, leukotrienes have been recognized as potent inflammatory mediators that initiate and propagate a diverse array of biologic responses including macrophage activation, mast cell cytokine secretion and dendritic cell maturation and migration [35, 36]. Thus, it is likely that LTs could play an important role in the hyperimmune/inflammatory storm observed in COVID-19 [29]. It has been proposed recently that montelukast, a leukotriene receptor antagonist, and zileuton, a 5-lipoxygenase inhibitor, might be possible treatment options for mild or even severe stages of COVID-19 [25-29], especially when given in combination [27].

Ze 339, a lipophilic subcritical CO_2_-extract prepared from the leaves of *Petasites hybridus* (L.) P.G. Gaertn., B Mey., & Scherb (Asteraceae), is an herbal treatment licensed in Switzerland and other countries used to treat allergic rhinitis [37]. Ze 339 has been demonstrated to inhibit leukotriene synthesis in various *in vitro* and *ex vivo* studies; the inhibition was solely attributable to the sum of petasins [38-41]. In particular, in human macrophages activated with platelet-activating factor (PAF), Ze 339 blocked Cys-LT and LTB4 synthesis, and further decreased PAF-and complement peptide C5a-mediated Cys-LT synthesis in eosinophils and LTB4 synthesis in neutrophils. In human eosinophils and neutrophils, the effects of the positive control zileuton, an orally active inhibitor of LT production, were similar to Ze 339 [40]. Furthermore, petasin and its isomers ehxibited no differences in their ability to inhibit 5-lipoxygenase (5-LOX), leukotriene C4 (LTC4) synthase and leukotriene A4 (LTA4) hydrolase, according to a recent study [38]. The authors of this study also showed that the extract matrix had no influence on the leukotriene inhibitory effects of the petasins. In addition to these mechanistic *in vitro* data, leukotriene levels decreased significantly in nasal lavage fluids of patients suffering from allergic rhinitis after 5 days of oral treatment with Ze 339 [39]. It is noteworthy to mention that Ze 339 also inhibited pro-inflammatory cytokine and chemokine response in human nasal epithelial cells after stimulation with viral mimics [42]. The authors also showed that the pro-inflammatory cytokine/chemokine response to bacteria was not inhibited by Ze 339. These findings highlight the potential of Ze 339 as a promising candidate for the treatment of a virally induced exacerbation of inflammatory processes in the upper airways. Based on the findings mentioned above, it was the aim of the present study to investigate if Ze 339 would also have an impact on SARS-CoV-2 replication *in vitro*. A cellular infection system was used to assess the ability of the substance(s) to interfere with cellular responses to and infection with SARS-CoV-2 virus variants.

## 2. Materials and Methods

### 2.1 Compounds and Extract

Ze 339 is a subcritical CO_2_-extract prepared from the leaves of *P. hybridus* (drug-extract ratio 50-100:1) and was manufactured by Max Zeller Söhne AG, Switzerland using a patented procedure [43, 44]. One coated film tablet contains 17-40 mg of Ze 339 and is standardized to 8 mg petasins. The herein used Ze 339 batch 150056 contained 37.7% total petasins and 27.2% fatty acids. The remaining 35.1% contained other constituents such as essential oils, sterols, minerals and vitamins. Pyrrolizidine-alkaloids have been quantitatively removed in the manufacturing process using an online-adsorption-technology and were no longer detectable (limit of quantification <2 ppb). A characteristic GC-chromatogram is shown in Supporting **Figure S1**. The extract was dissolved in a mixture of DMSO/H_2_O (50:50); a stock solution of 2 mg/ml was prepared and further diluted with assay buffer (DMEM, 2% FBS, 1%PS) to the final concentrations between 0.001 – 20 μg/m. Petasin (purity 92.82%), neopetasin (purity 94.49%) and isopetasin (purity 96.23%) were purchased from HWI Pharma Services GmbH, Ruelzheim, Germany. Stock solutions of petasin, isopetasin and neopetasin (10 mg/ml, respectively) were prepared using a DMSO/H_2_O (90:10) mixture and further diluted with assay buffer (DMEM, 2% FBS, 1%PS) to reach the final concentrations for testing (0.001 - 20 μg/ml, corresponding to 3.16 nM – 63 μM). Remdesivir (purity ≥98%) was purchased from Adipogen AG, Liestal, Switzerland. A 500 μM stock solution was prepared and further diluted with assay buffer to the desired concentrations (0.002-50 μM). To avoid cytotoxicity, the final DMSO concentration in all cellular experiments did not exceed 0.5%.

### 2.2 Cells

Vero E6 cells had been provided by the group of V. Thiel, Berne, Switzerland. Cells were cultivated in DMEM, high glucose, media (Gibco, Thermofisher AG, Allschwil, CH) and supplemented with Pen/Strep (1%, Bioconcept, Allschwil, CH) and 2% fetal bovine serum (FBS, Gibco, Thermofisher), at 37°C in a humidified atmosphere with 5% CO_2_.

### 2.3 Viral reconstitution

Virus stocks of the initial Wuhan strain of SARS-CoV-2 were provided by G. Kochs, University of Freiburg, D, (SARS-CoV_FR-3) and by EVAglobal virus archive, (SARS-CoV-2 strain /NL/2020 -AMS). Virus stocks were propagated in a Biosafety level 3 facility by infecting Vero E6 cells at a multiplicity of 0.1 and harvesting culture supernatant on day 3. Cell-free virus in culture supernatant was quantified by RT-PCR of the S-gene RDB region [45] and by plaque titration. Viral titers were determined by plaque assays in Vero cells.

### 2.4 Plaque Assay Protocol

Antiviral activity was determined by the degree of inhibition of viral cytopathic effect (CPE). Briefly, Vero E6 cells were seeded at a density of 3×10^6^ cells/96-well plate (ca. 3×10^4^ per well) one day before infection. On the day of infection, cells (ca. 80% confluent) were incubated with the compound of interest in the above-mentioned concentration range to a layer of uninfected cells. After 2h of incubating cells with the compound at 37°C in a humidified CO_2_ atmosphere, cells were infected with the virus with 100 plaque-forming units (pfu) per well to assess inhibitory effects on virus propagation. One hour post viral infection, cultures were overlayed with 100 μl low-melting agarose. Agarose (Bio-Rad Europe GmbH, Basel, CH), in DMEM/2% FBS was heated to melt and then cooled in a waterbath to a temperature of <40°C, and used to overlay cells in the pre-seeded, infected culture. Then, cells were incubated at 37 °C for approx. 48 h, within which a virus-driven CPE plaque formation is routinely observed in untreated controls. For virus inactivation, 50 μl of formaldehyde (15% w/v) were added for 10 minutes to the cultures without removing the low-melting agarose. After this period, fixative, culture medium and agarose were aspirated, and crystal violet (0.1% w/v) (Sigma-Aldrich) was added to each well and incubated for 5 min. Afterwards, the fixed and stained plates were gently rinsed several times under tap water and dried prior to enumeration. Antiviral activity was determined by the degree of inhibition of viral cytopathic effect (CPE), which became apparent in the form of distinct plaques forming in the cell layer. As the number of plaques observed per well was not easily distinguishable by eye, plates were scanned, and the images were counted as described by Honko et al. [46] using Image J 1.53k software [47]. For image processing, images were made binary images according to the Image J definition, the limit to threshold option was enabled under ‘set measurements’ and pixels of the selected area were counted. The results were normalized to positive (virus infected) and negative (DMSO control) controls in each assay plate.

### 2.5 Cytotoxicity Testing

Vero E6 cells were plated as described above, 3×10^6^/well in 100 μl complete culture medium/2% FBS and cultured overnight. The drug of interest (stock solution) was diluted in DMEM supplemented with 2% FBS, and 50 μl was then added per well (in duplicates or triplicates at the indicated final concentration). The plates were cultured for 48 h, similar to the duration of the infections, after which they were fixed and stained with crystal violet as described above. Cytotoxic effects were evaluated using Image J as described above.

### 2.6 Statistics

Compounds were tested in duplicates or triplicates per experiment, each experiment was repeated at least twice. Antiviral data were fitted to a sigmoidal curve and a four parameter logistic model was used to calculate IC_50_ values using the equation: Y=Bottom + (Top-Bottom)/(1+10^((LogIC50-X)*HillSlope)). The IC_50_ values are reported at 95% confidence intervals. This analysis was performed using GraphPad Prism v.9.2.0 (San Diego, USA).

## 3. Results

### 3.1. Inhibition Assay of Remdesivir, Ze 339, Petasin, Isopetasin and Neopetasin against the original SARS-CoV-2 Wuhan variant

In this study, the potent antiviral effects of the *Petasites hybridus* CO_2_-extract Ze 339 as well as its’ active compounds isopetasin and neopetasin (**Figure 1**) against SARS-CoV-2 infection of the Vero E6 cell line (derived from primary embryonic monkey kidney epithelial cells) was demonstrated.

**Figure 1:**
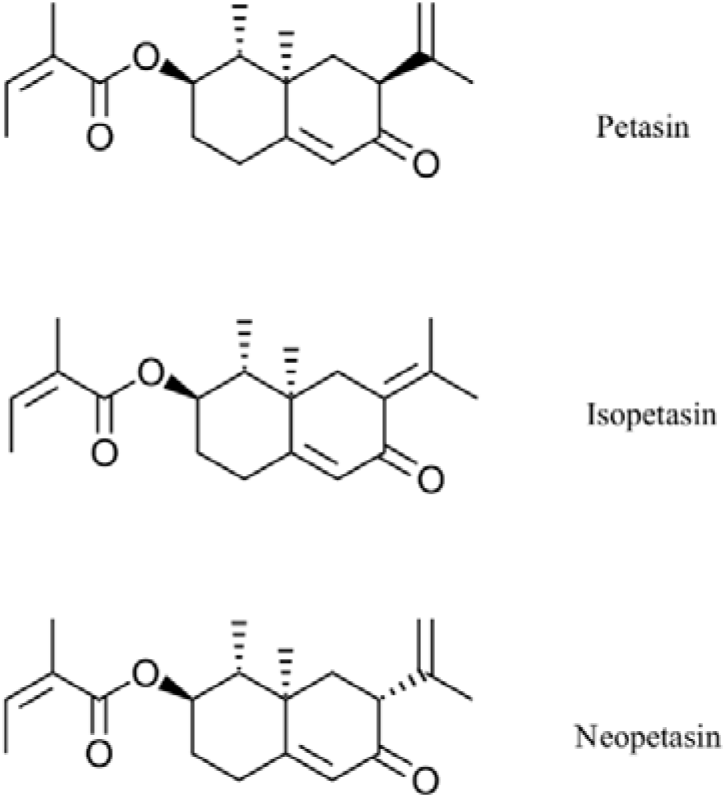
Chemical structures of the main petasins from *Petasites hybridus*

The antiviral activity of the test compounds was assessed as previously described [46, 48] by visualization of the extent of the cytopathogenic effect (CPE) in the form of cytolytic plaque formation on Vero E6 cells when infected with a clinical strain of SARS-CoV-2, Wuhan. Vero E6 cells are stable cell lines expressing a high level of the ACE2 receptor [49]. They were recently employed in studies to assess SARS-CoV-2 infection and replication by quantifying the virally induced CPE leading to the formation of cytolytic plaques [50-53]. Prior to antiviral testing the cytotoxicity of all samples towards Vero E6 cells was assessed, and the 50% cytotoxic concentration (CC_50_) of the test compounds was determined. In addition, a selectivity index (SI) was determined as CC_50_/IC_50_.

As shown in **Figure 2 & 3**, Ze 339 as well as petasin, isopetasin and neopetasin did not show any cytotoxic effect in this assay format up to concentrations of 20 μg/ml (63.2 μM for the petasines, respectively).

**Figure 2:**
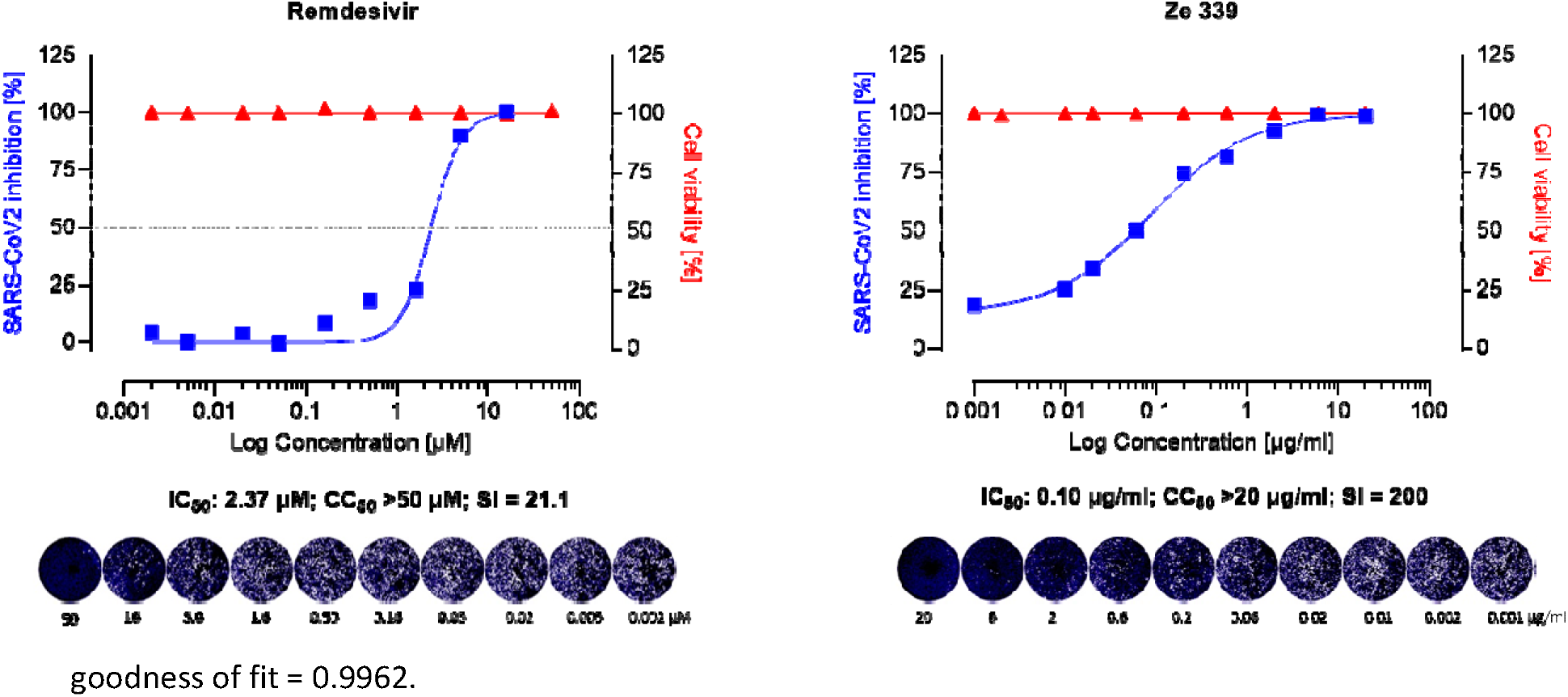
Anti-SARS-CoV-2 activity of the remdesivir and Ze 339 in VeroE6 cells was assessed using the Wuhan strain of SARS-CoV-2. Cells were infected with 100 pfu of a clinical virus isolate in the presence of the indicated concentrations. Blue squares represent inhibition of SARS-CoV-2 infection (%), red triangles represent cell viability (%). Data are expressed as the mean of two replicates (infection performed in duplicate) with their respected 95% confidence interval. Remdesivir 95% confidence interval = 0.25-0.51, goodness of fit = 0.9668; Ze 339 95% confidence interval = 0.06-0.12, goodness of fit = 0.9962.

To investigate whether the *P. hybridus* CO_2_-extract Ze 339 and its main active components petasin, isopetasin and neopetasin exert antiviral activity, Vero E6 cells, infected with the Wuhan variant of SARS-CoV-2 were incubated for 48h in the presence of Ze 339 (0.001-20 μg/ml) or the petasines (0.001-20 μg/ml, corresponding to 0.0031-63.2 μM), respectively. The results demonstrated that Ze 339 (**Figure 2**) inhibited the formation of virus-driven cytopathic changes in a dose-dependent manner in infected Vero E6 cells with an IC_50_ of 0.12 μg/ml (C.I. 0.06-0.12). The antiviral drug remdesivir was used as reference and showed anti-SARS-CoV-2 activity with an IC_50_ of 2.37 μM (C.I. 0.25-0.51) (**Figure 2**).

Similar IC_50_ values ranging between 1-11 μM for remdesivir have been reported in the literature [51, 52, 54]. Compared to remdesivir, isopetasin and neopetasin also exerted potent antiviral activities with IC_50_ values ranging from 0.79 μM – 1.2 μM (**Figure 3**). For petasin an IC_50_ of 10.79 μM was determined, which was higher than that of isopetasin and neopetasin. However, in aqueous solution petasin converts quickly to its more stable stereoisomer isopetasin. According to a recent screening of 5,632 compounds (natural products as well as synthetics), including 3,488 compounds that have undergone clinical stage testing across 600 indications, only 19 compounds were identified as having an IC_50_ in the nanomolar (<1 μM) range, when tested against SARS-CoV-2 in Caco-2 cells [55]. Among those compounds tested were nafamostat and camostat which have different mechanisms of antiviral action [55]. The potent anti-SARS-CoV-2 activity of isopetasin and neopetasin is therefore promising. However, further steps are necessary to evaluate the precise underlying mechanism of the antiviral action.

**Figure 3:**
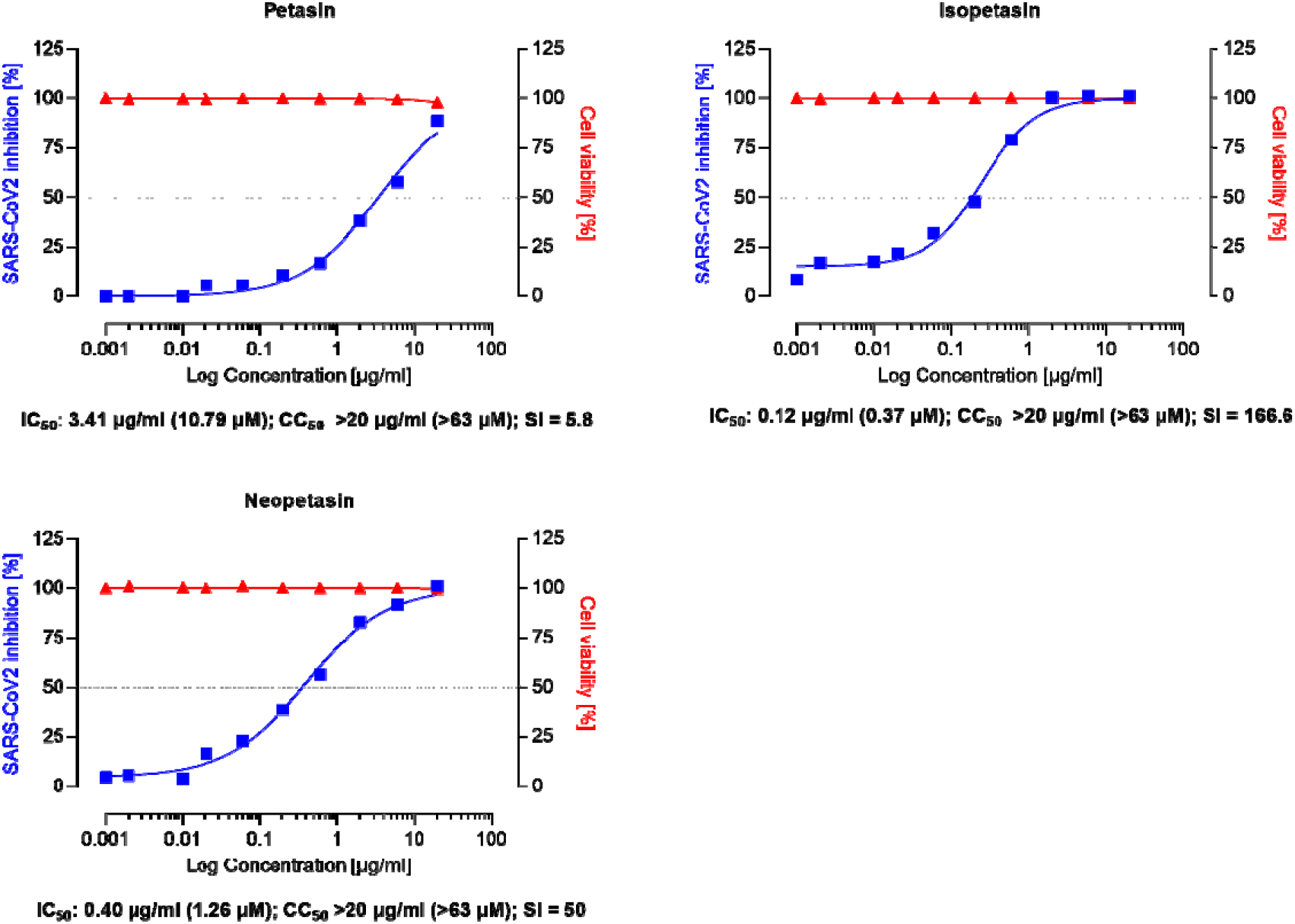
Anti-SARS-CoV-2 activity of the isopetasin and neopetasin in VeroE6 cells was assessed using the Wuhan strain of SARS-CoV-2. Cells were infected with 100 pfu of a clinical virus isolate in the presence of the indicated concentrations. Blue squares represent inhibition of SARS-CoV-2 infection (%), red triangles represent cell viability (%). Data are expressed as the mean of two replicates (performed in duplicate) with their respected 95% confidence interval. Petasin confidence interval = 2.70-4.31, goodness of fit = 0.9874; Isopetasin 95% confidence interval = 0.15-0.41, goodness of fit = 0.9918; neopetasin 95% confidence interval = 0.27-0.55, goodness of fit = 0.9929.

The selectivity index (SI) of a compound is a commonly used metric to express a compound’s *in vitro* efficacy in inhibiting virus reproduction and an important measure to compare the antiviral efficacy of experimental drugs [56]. The greater the SI ratio, the more successful and safe a medicine should be for treating a viral infection *in vivo*. The ideal medicine would be cytotoxic at high doses but antiviral at very low concentrations, resulting in a high SI value and the ability to remove the target virus at concentrations much below its cytotoxic concentration. In our study, Ze 339 showed an SI of 200, compared to an SI of 21.1 for remdesivir; the SI for Isopetasin is 80 and for neopetasin 50 when tested against the Wuhan virus variant. Petasin had the lowest SI value but as mentioned above, the values for petasin are likely to be affected by a rapid isomerization. It is noteworthy to mention that compared to remdesivir, Ze 339 as well as isopetasin and neopetasin exerted a higher selectivity index. This is in so far of importance since a vast majority of drug candidates, including repurposed drugs, being currently evaluated as COVID-19 treatments, have SIs that are much lower than that achieved by Ze 339 or the petasines [57].

### 3.1. Inhibition Assay of Remdesivir, Ze 339 and Isopetasin against the SARS-CoV-2 Delta variant

As novel SARS-CoV-2 variants of clinical concern spontaneously develop around the world, we tested the antiviral activity of Ze 339 and isopetasin also against the currently dominant SARS-CoV-2 Delta variant. In a first proof of concept experiment, the effect of Ze 339 as well as isopetasin were investigated in the plaque reduction assay when added to the cells prior to infection with the Delta variant. For this experiment, only isopetasin was chosen of the petasins since it is the most stable isomer and since it exerted the lowest IC_50_ value when tested against the Wuhan variant. After infection of Vero E6 cells with the Delta virus variant, Ze 339 showed a slightly lower antiviral activity with an IC_50_ of 0.40 μg/ml and isopetasin with an IC_50_ of 0.28 μg/ml (0.88 μM) (**Figure 4**). A comparable SARS-CoV-2 inhibitory activity of isopetasin and remdesivir against the Delta variant was observed in this infection experiment. When compared to the Wuhan variant, the activity of Ze 339 and isopetasin against the Delta variant were comparable.

**Figure 4:**
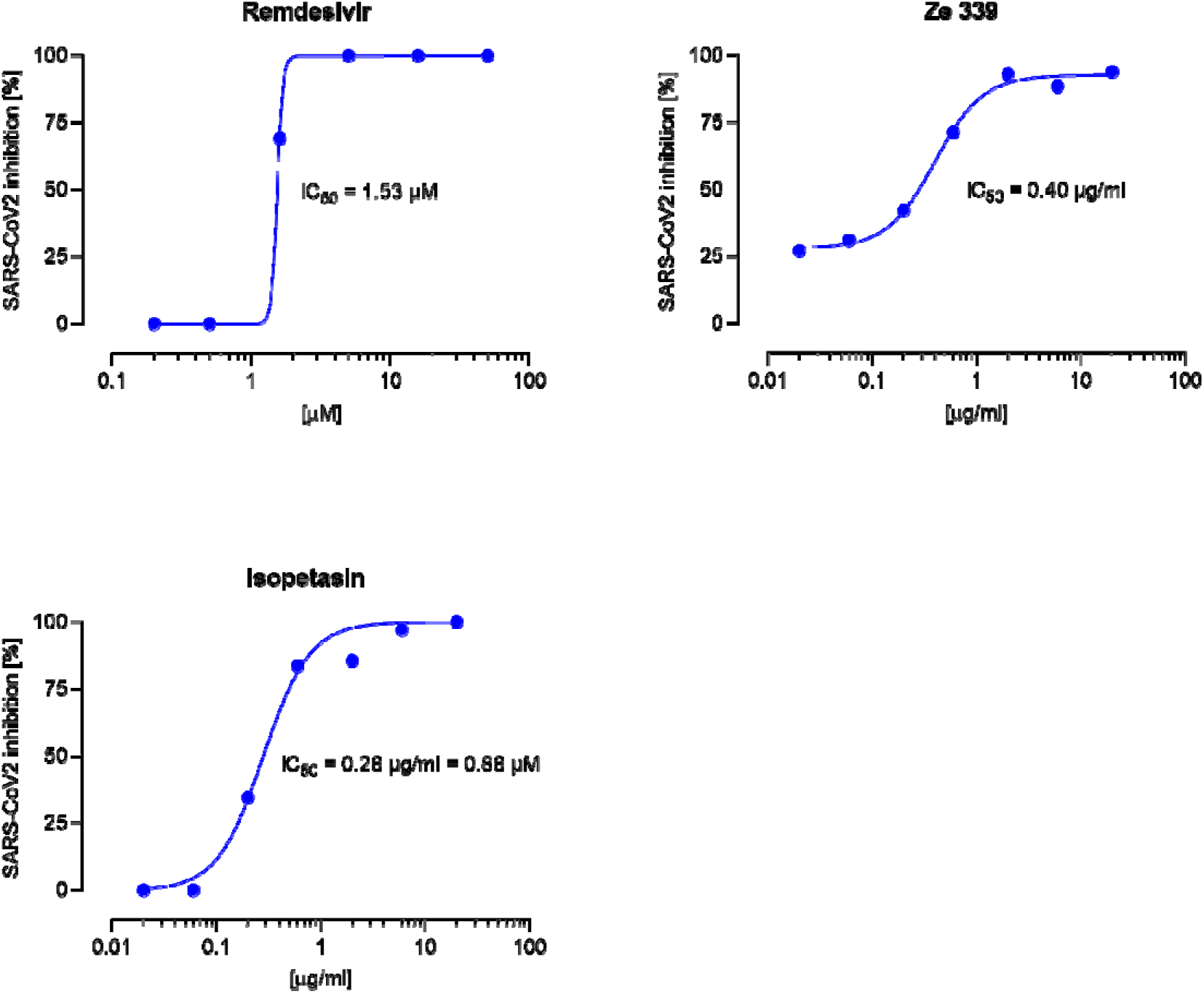
Dose response curves for remdesivir, Ze 339, and isopetasin on the SARS-CoV-2 Delta variant. Cells were infected with 100 pfu of a clinical virus isolate in the presence of the indicated concentrations. Blue circles represent inhibition of SARS-CoV-2 infection (%). Data are expressed as the mean of duplicate measurements with their respected 95% confidence interval. Ze 339 95% confidence interval 0.26-0.58, goodness of fit = 0.9943, isopetasin 95% confidence interval = 0.21-0.38, goodness of fit = 0.9845.

## 4. Discussion

Facing a globally rapidly expanding new disease such as COVID-19, drug repurposing of existing moieties appears to be a very favorable approach as their development process to drug approval can utilize pre-existing preclinical and often clinical or post-marketing data [55, 58].

The current study demonstrates that Ze 339, isopetasin as well as neopetasin were able to potently protect cells from SARS-CoV-2 infection. The antiviral activity of Ze 339 (IC_50_ 0.10 μg/ml) against the initial Wuhan strain clearly exceeded that of remdesivir (IC_50_ 1.42 μg/ml). Similar potencies were observed for the Delta variant, were Ze 339 protected cells from an SARS-CoV-2_D infection with an IC_50_ of 0.40 μg/ml (remdesivir: IC_50_ of 0.91 μg/ml). The experiments with isolated petasins suggest that the antiviral effect of Ze 339 is due to the sum of petasins.

Easy to use and affordable COVID-19 therapies are urgently needed for clinical disease management. A great advantage of Ze 339 therefore is that it already is an approved OTC drug in Switzerland and various other countries for oral treatment of seasonal allergic rhinitis. Further, Ze 339 is available for self-medication. Clinical and postmarketing surveillance studies did not reveal any safety concerns [59-63]. The range and incidence of adverse effects in the respective studies were very low; described adverse events had all previously been known and are included in the summary of product characteristics [64]. Furthermore, as an approved drug, Ze 339 is subject to pharmacovigilance activities according to current standard procedures. The lack of safety findings reported for *P. hybridus* extract Ze 339 reflects the fairly robust overall safety assessment from clinical experience exceeding 15 years since marketing authorization.

In addition, the mechanism of action as well as the active compounds of Ze 339 have been well-described. From a range of pharmacological studies, Ze 339 and its active constituents, petasin and its isomers iso- and neopetasin, have been shown to ultimately inhibit the synthesis of leukotrienes in human target cells such as leukocytes and macrophages cultured *in vitro* or *ex vivo* [39-41, 65]. Moreover, Ze 339 has been shown to reduce TNF-α, IL-6 and IL-8 levels in human nasal epithelial cells challenged with viral mimetics [42].

Furthermore, in addition to the above mentioned mechanisms of action, petasin is also a potent activator of the AMP-activated protein kinase (AMPK) [66]. AMPK impairs SARS-CoV-2 replication in several ways: activated AMPK phosphorylates angiotensin-converting enzyme 2 and decreases SARS-CoV-2 binding to the enzyme, thereby decreasing cellular uptake of the virus [67]. Activated AMPK also suppresses the protein kinase B (AKT)/mechanistic target of rapamycin (mTOR) pathway, which is required for viral protein translation and replication [68]. This is consistent with previous clinical data showing an improved mortality in diabetic COVID-19 patients trerated with the indirekt AMPK activator metformin [69-71]. Finally, metformin suppresses inflammatory pathways and immunological responses (cytokine storms) [72].

It is further worth mentioning that the mechanism of anti-inflammatory action of Ze 339 is comparable to that of montelukast and zileuton, which are currently under discussion as therapeutic candidates for the treatment of COVID-19 [25-29]. In a recent study by Durdagi et al. [73] demonstrated the neutralization effect of montelukast on SARS-CoV-2 *in vitro* by showing that virus replication could be significantly delayed. The authors suggested to consider montelukast for prophylactic treatment. In our current study, a pre-incubation of cells with the test compounds for 2 h prior to virus infection had a clear positive effect, suggesting a possible prophylactic role also for the petasins. While the montelukast IC_50_ values were at 18.82 μM and 25 μM in *in vitro* cell culture models of SARS-CoV-2 activity [73, 74], the IC_50_s for Ze 339, isopetasin and neopetasin determined in the current study were significantly lower. The visibly shallower inhibition curves of the petasin compounds compared to the antiviral mechanism of remdesivir could suggest a more complex mode of interference with the viral life cycle which is supported by the above mentioned various mechanisms of action of Ze 339. A similar difference is also been observed for different HIV inhibitor classes (e.g. nucleosidic polymerase inhibitors versus protease inhibitors; not shown).

## 5. Conclusions

In conclusion, since affordable therapies for treating SARS-CoV-2 infection and COVID-19 are urgently needed, we propose that Ze 339 could be a promising candidate for drug repurposing with a supposed dual mode of action through a direct anti-SARS-CoV2 effect and a potent indirect inhibitory effect on leukotriene biosynthesis/cytokine activity or AMPK activation.

Taken together, the *Petasites hybridus* subcritical CO_2_-extract Ze 339 may provide a safe, low-cost alternative for treating patients infected with SARS CoV-2. As Ze 339 is registered in Switzerland as OTC drug, it is a promising repurposing candidate due to its proven safety profile. For further development, the current *in vitro* data will have to be verified *in vivo*, and a clinical benefit of Ze 339 as a possible drug for treating mild to moderate cases of COVID-19 needs to be proven in future clinical studies.

## Supplementary Materials

**Figure S1**: A GC chromatogram of Ze 339 is shown in **Figure S1**.

## Author Contributions

The following statements should be used “Conceptualization, T.K., M.K., V.B. and J.D; methodology, L.U.; software L.U., V.B.; formal analysis, V.B.; G.B. writing—original draft preparation, V.B.; G.B., T.K. writing—review and editing, J.D.; visualization, V.B.; supervision, T.K.;. All authors have read and agreed to the published version of the manuscript.”

## Data Availability Statement

Data are available on request from the authors.

## Acknowledgement

For providing access to the SARS-CoV-2 strain /NL/2020 – AMS, we thank the EVA GLOBAL consortium (funded by the European Union’s Horizon 2020 research and innovation programme under grant agreement No 871029) and Dr. B. Hagman.

## Supplemental material to

**Figure S1:**
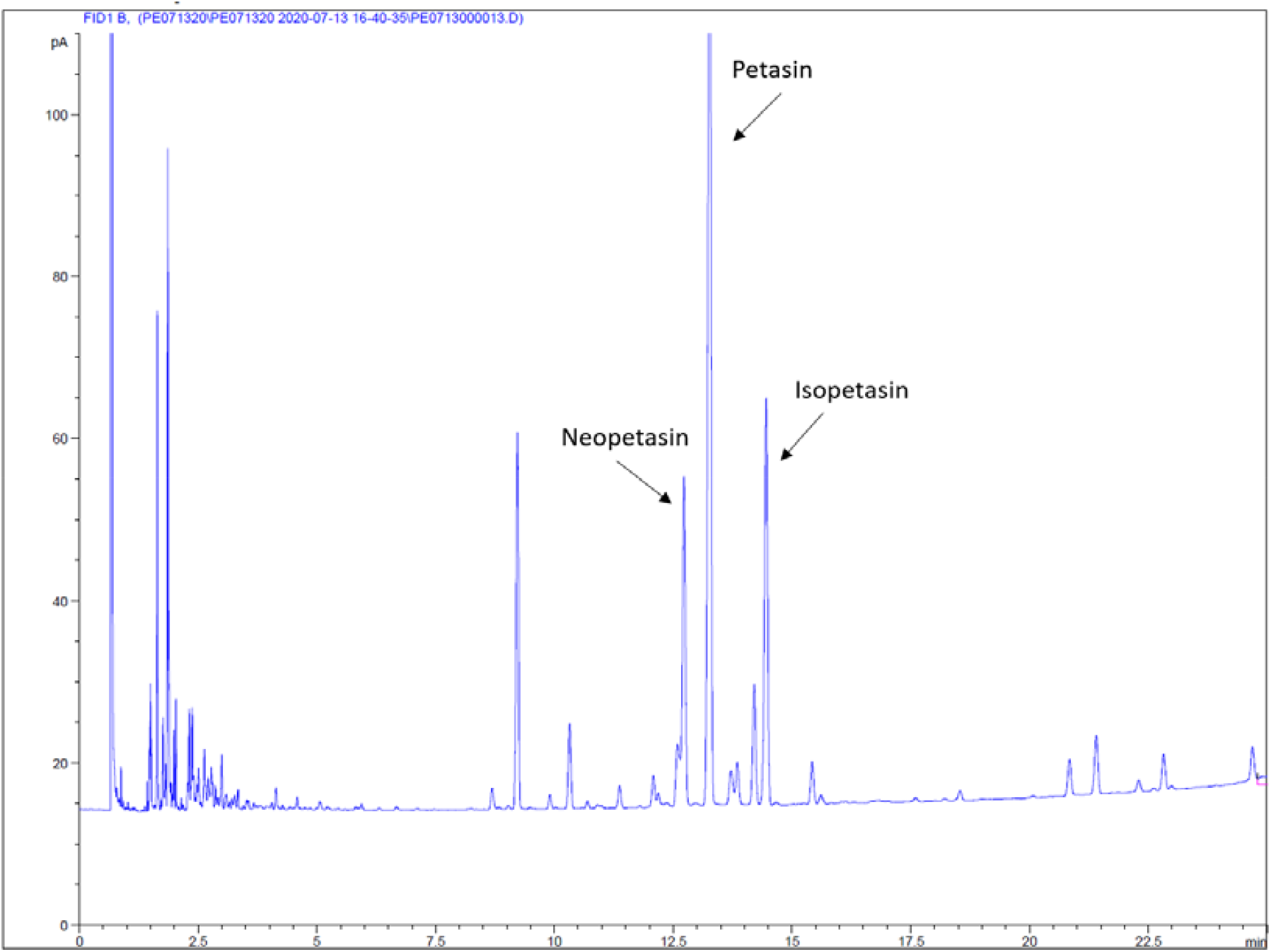
Gas chromatogram of *Petasites hybridus* leaf extract Ze 339 (batch 150056). Quantitative determination of total petasins (petasin, isopetasin, neopetasin), the active compounds of Ze 339 using gas chromatography and a flame ionization detector (FID). GC-column 100% polydimethylsiloxane (e.g. DB-1, length: 25 m, ID: 0.32 mm, dF: 0.52 μm); Injector temperature: 270 °C; Injection volume 1 μL.

